# Selection on phenotypic plasticity favors thermal canalization

**DOI:** 10.1101/2020.06.11.146043

**Authors:** Erik I. Svensson, Miguel Gomez-Llano, John T. Waller

## Abstract

Climate change affects organisms worldwide with profound ecological and evolutionary consequences, often increasing population extinction risk. Climatic factors can increase the strength, variability or direction of natural selection on phenotypic traits, potentially driving adaptive evolution. Phenotypic plasticity in relation to temperature can allow organisms to maintain fitness in response to increasing temperatures, thereby “buying time” for subsequent genetic adaptation and promoting evolutionary rescue. Although many studies have shown that organisms respond plastically to increasing temperatures, it is unclear if such thermal plasticity is adaptive. Moreover, we know little about how natural and sexual selection operate on thermal reaction norms reflecting such plasticity. Here, we investigate how natural and sexual selection shape phenotypic plasticity in two congeneric and phenotypically similar sympatric insect species. We show that the thermal optima for longevity and mating success differ, suggesting temperature-dependent trade-offs between survival and reproduction. Males in these species have similar thermal reaction norm slopes but have diverged in baseline body temperature (intercepts), being higher for the more northern species. Natural selection favoured reduced thermal reaction norm slopes at high ambient temperatures, suggesting that the current level of thermal plasticity is maladaptive in the context of anthropogenic climate change and that selection now promotes thermal canalization and robustness. Our results show that ectothermic animals also at high latitudes can suffer from overheating and challenge the common view of phenotypic plasticity as being beneficial in harsh and novel environments.

**Significance Statement:** Organisms are increasingly challenged by increasing temperatures due to climate change. In insects, body temperatures are strongly affected by ambient temperatures, and insects are therefore expected to suffer increasingly from heat stress, potentially reducing survival and reproductive success leading to elevated extinction risks. We investigated how ambient temperature affected fitness in two insect species in the temperate zone. Male and female survivorship benefitted more from low temperatures than did reproductive success, which increased with higher temperatures, revealing a thermal conflict between fitness components. Male body temperature plasticity reduced survival, and natural and sexual selection operated on such thermal plasticity. Our results reveal the negative consequences of thermal plasticity and show that these insects have limited ability to buffer heat stress.

## Introduction

Ongoing climate change due to anthropogenic activities affects organisms across the globe, often causing local extinctions(1–3) and changing selection pressures on phenotypic traits(4, 5). Ectothermic animals such as reptiles, amphibians and arthropods are especially sensitive to climate change, as their body temperatures are strongly influenced by ambient temperatures, with strong effects on life-history traits such as growth, survival and reproduction(6). Sensitivity to ambient temperatures is thought to be especially high in low-altitude tropical ectotherms living in shaded habitats such as forests, as these organisms are adapted to rather stable and narrow thermal zones(7–9). When critical thermal limits are exceeded in such organisms(10), extinction risk is thought to increase dramatically(2, 11). Recent studies have shown that upper critical thermal limits are strongly conserved phylogenetically(12) but also that ectothermic animals at high latitudes can experience overheating(10).

Increasing local, regional and global temperatures could therefore elevate population extinction risk, but an alternative outcome is that natural or sexual selection(5, 13) instead promote thermal adaptation(6). This could increase the ability to withstand high temperatures, resulting in evolutionary rescue(14). Evolutionary rescue through genetic adaptation would take time, due to a delay between the onset of selection and genetic evolutionary responses. However, phenotypic plasticity (environment-dependent expression of alternative phenotypes by the same genotype) and thermal plasticity, in particular, could “buy time” for organisms before establishment of evolutionary adaptation by genetic means(15–18). Recent research on endothermic organisms like birds have revealed between-generation plastic changes in life-history traits in response to increasing temperatures, through earlier breeding phenologies(19, 20). However, the adaptive significance of such thermal plasticity is largely unknown and in some cases plasticity might be non-adaptive or even maladaptive(21–24). A recent review on the limited evidence for selection on thermal plasticity found only eight examples of selection studies on thermal plasticity, from four species of plants and two species of birds (in the latter case no direct measurements of temperature were used but indirect proxies such as year or climate index)(25). Notably, recent meta-analyses have not documented a single selection study on thermal plasticity in insects, the most species rich animal class and a group to be strongly affected by increasing temperatures(20, 25). A recent experimental evolution study in a laboratory setting suggested that sexual selection could facilitate evolutionary rescue in response to increasing temperatures(26). However, there is not yet any field study to our knowledge, on how natural and sexual selection operate on thermal plasticity and how such selection might promote thermal adaptation.

Here, we quantified fitness effects of temperature and natural and sexual selection on thermal plasticity in natural populations of two sexually dimorphic insect species: the banded demoiselle (*Calopteryx splendens*) and the beautiful demoiselle (*C. virgo*). These two closely related species belong to the odonate superfamily Calopterygoidea, which has a tropical origin in southeast Asia(27, 28). Wing pigmentation in these and other damselflies and dragonflies have previously been shown to be important in intra- and intersexual selection as well as in speciation and species recognition and does also play a role in thermal adaptation(13, 27, 29). The tropical origin of the genus *Calopteryx* would presumably make these species well adapted to high temperatures and sensitive to low temperatures. Yet, these two species are the only members of this clade that have successfully expanded into northern Europe and Scandinavia, where this study took place. Results in this study are also of general for those who wish to understand sex differences in the fitness effects of ambient temperatures and how sexual selection on males might increase thermal adaptation and contribute to evolutionary rescue (14, 30–32).

## Results

Both males and females in these temperature-sensitive insects are strongly affected by ambient temperatures under field conditions, in terms of both survivorship and mating success (Fig. 1A-H; Tables S1-S2). Longevity showed a curvilinear relationship with temperature, and typically peaked between 15 and 20 °C for males and females in both *C. virgo* and *C. splendens* (Figs. 1A, B, E, F,Table S1-S2). Mating success also showed significant curvilinear relationship with temperature for all phenotypes, except *C. virgo* females (Fig. 1C, D, G, H; Tables S1-S2). In contrast to longevity where the thermal optima were found between 15 and 20 °C, the fitness peaks for mating success were located between 20 and 25 °C for *C. virgo* males (Fig. 1C) and between 25 and 30 °C for the other three phenotypes (Fig. 1D,G,H). Thus, the thermal optima differed for longevity and mating success, with mating success benefitting from higher temperatures whereas survivorship being negatively affected (Fig. 1A-H).

**Fig. 1.**
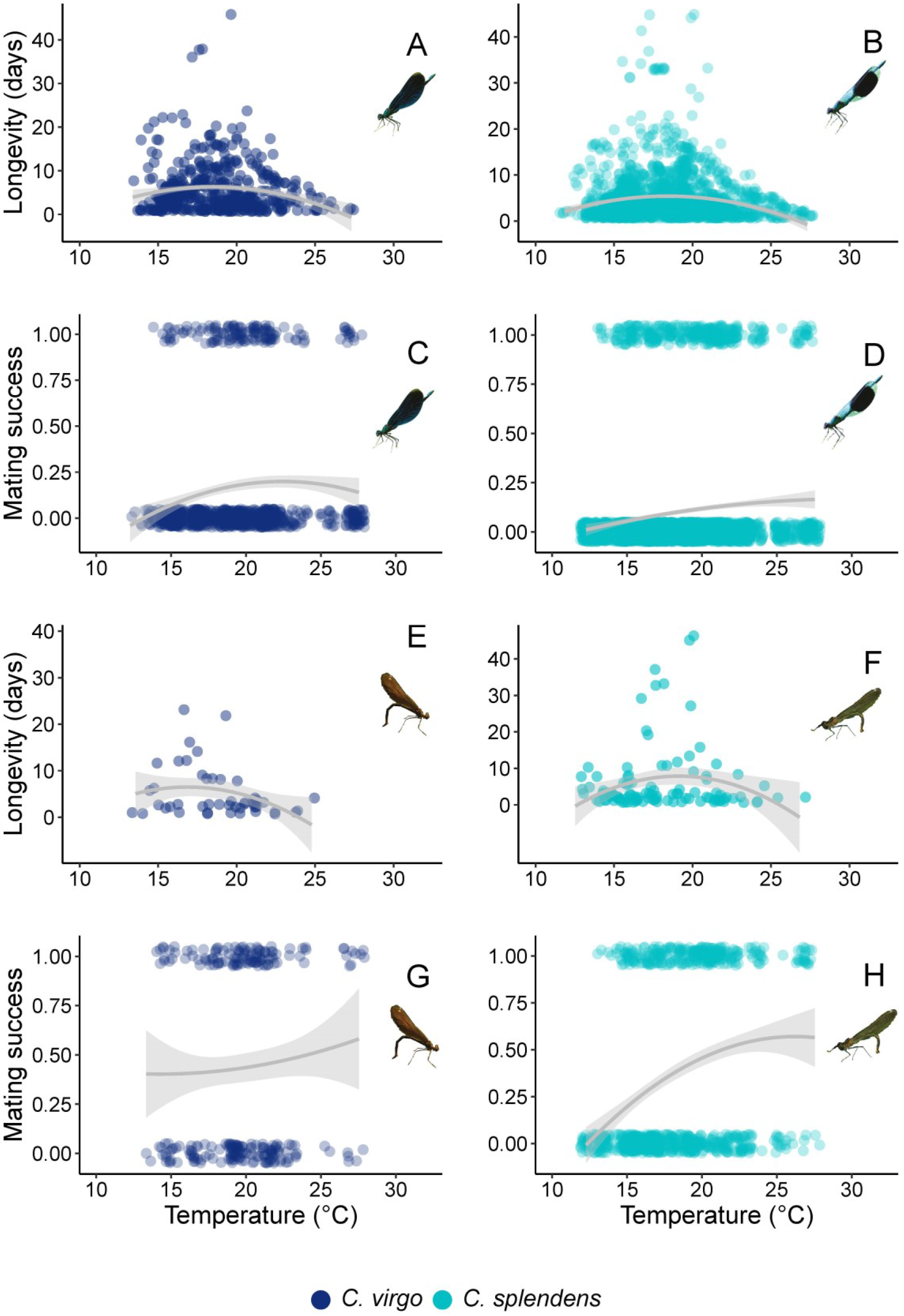
Effects of ambient temperatures on fitness components in male and female *Calopteryx* damselflies. We quantified how survivorship (longevity) and mating success was affected by ambient temperature in a natural field population (Klingavälsån at Sövdemölla in Southern Sweden) along a slowly running stream. Inserted are pictures of the four phenotypes: males and females of *Calopteryx virgo* (left column) and males and females of *C. splendens* (right column). **A:** Longevity in *C. virgo* males. **B:** Longevity in *C. splendens* males. **C:** Mating success in *C. virgo* males. **D:** Mating success in *C. splendens* males. **E:** Longevity in *C. virgo* females. **F:** Longevity in *C. splendens* females. **G:** Mating success in *C. virgo* females. **H:** Mating success in *C. splendens* females. See Tables S1 and S2 for statistical tests.

To investigate how natural and sexual selection operate on thermal plasticity and thermal reaction (6) in males, we used thermal imaging, a novel non-invasive tool to quantify body surface temperatures in insects and other animals(27). Thermal imaging estimates surface temperatures of animals(33). For small organisms like insects, with their low thermal inertia, surface temperatures are highly correlated with internal body temperatures(27) (Fig. S1). We estimated variation between individual males of both species in heating rates of their thorax (containing the wing muscles, which are essential for flight) following a cold challenge (Fig. 2A). This experimental setup simulates the effects of a cold night, after which the males are gaining heat in the early morning hours before they start searching for females and food under natural conditions in the field. We estimated thermal plasticity as the slope of the thermal reaction norms (°C/second) and intercept (baseline temperature in °C) at the start of the experiment, controlling for the ambient temperature. Although, these two aspects of reaction norms (slope and intercepts) are the two classical parameters in theoretical models for the evolution of plasticity and canalization(6, 15, 16), selection on such parameters have seldom been quantified(25).

**Fig. 2.**
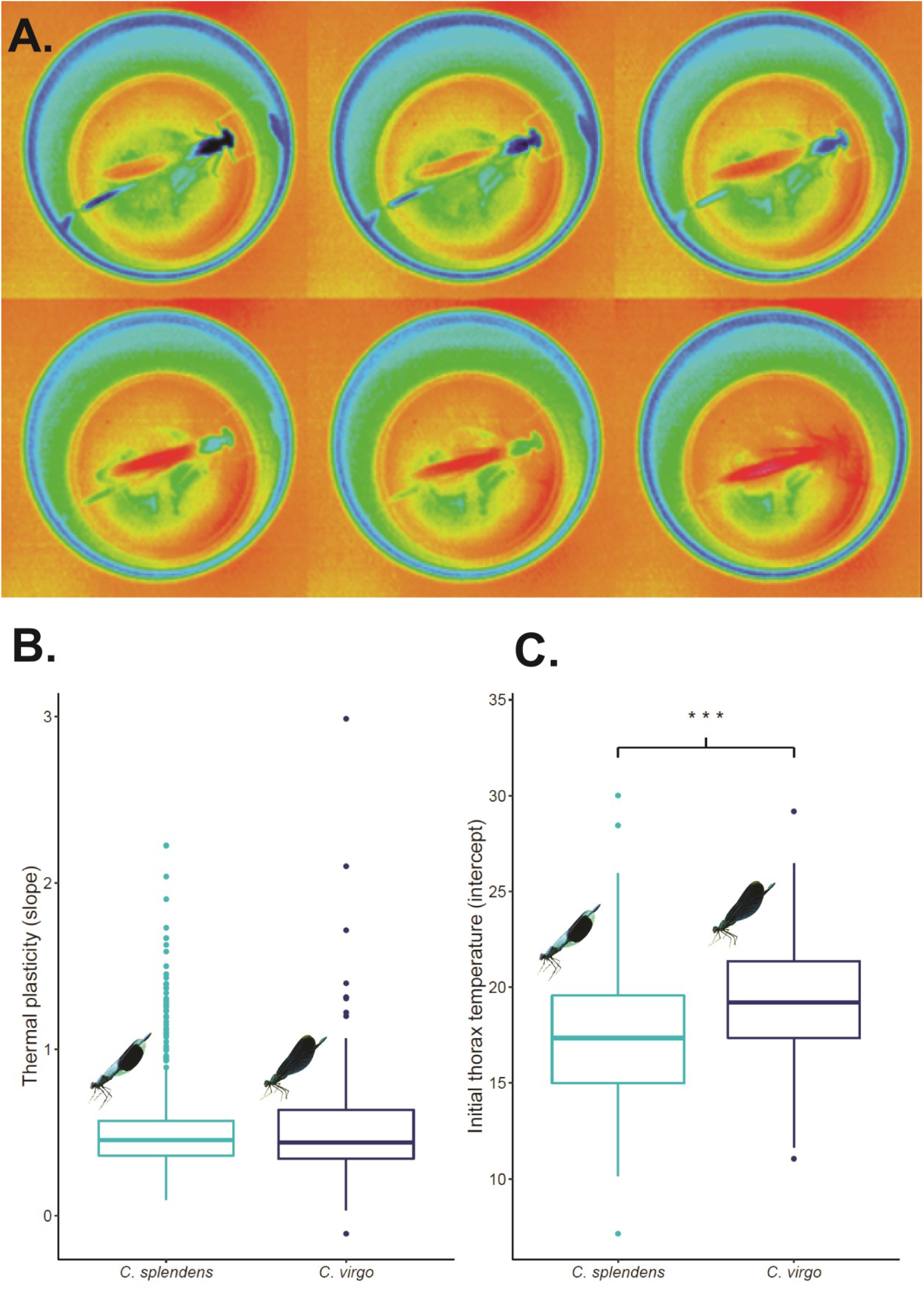
Using thermal imaging (infrared camera) to quantify species-specific thermal reaction norm slopes and intercept in male *Calopteryx* damselflies. **A:** We recorded individual variation in heating rates using thermal image camera of male *Calopteryx* damselflies in the field, to quantify variation in thermal reaction norms (slopes and intercepts). Shown is the same individual male that is heating up under ambient temperature, after being cooled down in a fridge box (c. a. 5 °C; Upper left shows first picture). Warmer colours (red, orange) show higher surface body temperature. We recorded a thermal image every 10^th^ second for three minutes until he flew off, which usually happened before the three minutes (180 seconds) were over. **B-C:** individual and species variation in thermal reaction norms of *C. splendens* (left; n = 778) and *C. virgo* (n = 179; see also Fig. S2). These two species do not differ in thermal reaction norm slopes (**B**), but they differ significantly in thermal reaction norm intercepts (**C**). Statistical tests refer to t-tests, using each individual-level reaction norm slopes and intercepts as individual data points (see Fig. S2 for range of variation). The higher intercept of *C. virgo* means that its baseline body temperature is higher at low ambient temperatures. This might explain its more northernly distributional range as it could become active at lower temperatures. See Table S3 for statistical tests.

We found that the slopes of the thermal reaction norms between these two closely related species were similar and almost identical (Fig. S2). Thus, thermal plasticity (slopes) are conserved between these two phylogenetically closely related congeneric species (Fig. 2B; Table S3). In contrast, the reaction norm intercept (baseline temperature) was significantly higher in *C. virgo* than in *C. splendens* (Fig. 2C; Table S3). Thus, the more melanised species *C. virgo* seemed to be more cold-tolerant and better able to keep a high body temperature at low ambient temperatures (Fig. 2C).

Next, we estimated natural and sexual selection on thermal reaction norm slopes and intercepts in males of these two species by combining data on individual thermal reaction norms with field data on male longevity and mating success. We found evidence for significant natural and sexual selection on thermal reaction norm slopes and intercepts in both species (Fig. 3; Table S4). In *C. splendens,* natural and sexual selection on reaction norm slopes and intercepts had opposite signs consistent with a trade-off, with longevity decreasing with increasing slopes and intercepts, whereas mating success increased (Fig. 3A). In *C. virgo*, this selective trade-off was less evident and natural and sexual selection were more concordant (Fig. 3B, Table S4). Natural selection operated on the thermal slopes and intercepts in both these species, but in different ways, revealing species-specific selection on thermal reaction norms (Fig. 3A-B; Table S4). In contrast, sexual selection favored higher reaction norm intercepts in both species but did not operate on reaction norm slopes (Fig. 3; Table S4).

**Fig. 3.**
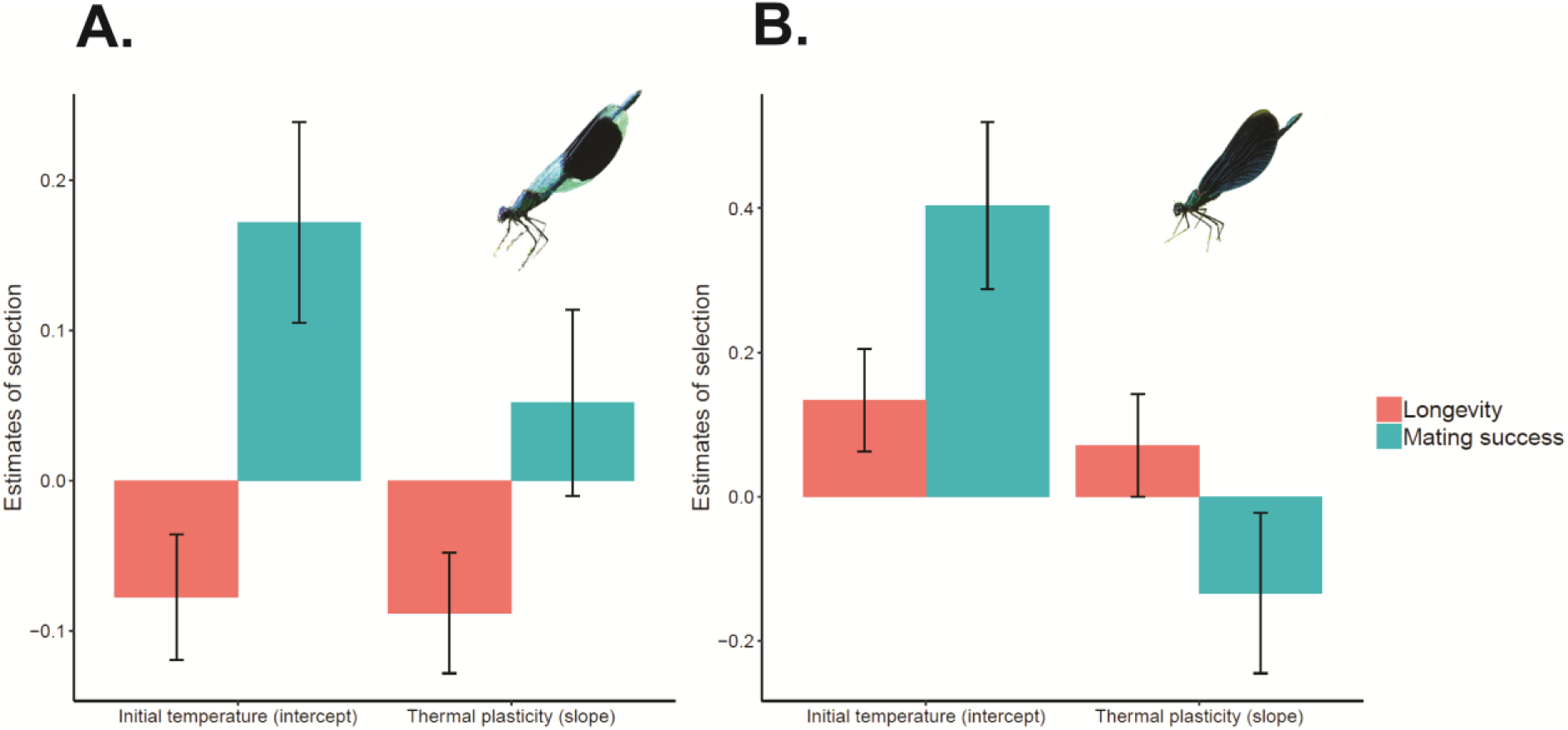
Natural and sexual selection on reaction norm slopes and intercepts in *Calopteryx* males across all seasons and ambient temperatures. Field estimates (standardized selection gradients) of natural selection (longevity) and sexual selection (mating success) on reaction norm slopes and intercepts across all years and temperature ranges for *C. splendens* males (longevity, n = 778; mating success, n = 2160) (**A**) and *C. virgo* males (longevity, n = 179; mating success, n = 413) (**B**). Shown is the standard error around the mean. See Table S4 for statistics.

Natural selection on thermal reaction norm slopes and intercepts were also strongly temperature-dependent in both species (Fig. 4A-F; Table S5). Specifically, all four interaction terms between ambient temperature and the thermal slopes and thermal intercepts were significant and negative in both species (Fig. 4A-D; Table S5). Thus, lower slopes (lower thermal plasticity) and lower intercept (lower baseline body temperature) were beneficial to survival at high ambient temperatures (Fig. 4A-D; Table S5). In contrast, at low and intermediate temperatures, higher slopes and higher intercepts were associated with high survival (Figs. 4A-D; Table S5). Hence, thermal plasticity was maladaptive when ambient temperature was high, and selection then instead favoured thermal canalization and lower body temperatures, presumably to avoid overheating (Figs. 4A-D; Table S5). Finally, mating success increased with higher thermal intercept in both species (Fig. 3; Table S4), and in *C. virgo* it increased with both temperature and intercept (Fig. 4E).

**Fig. 4.**
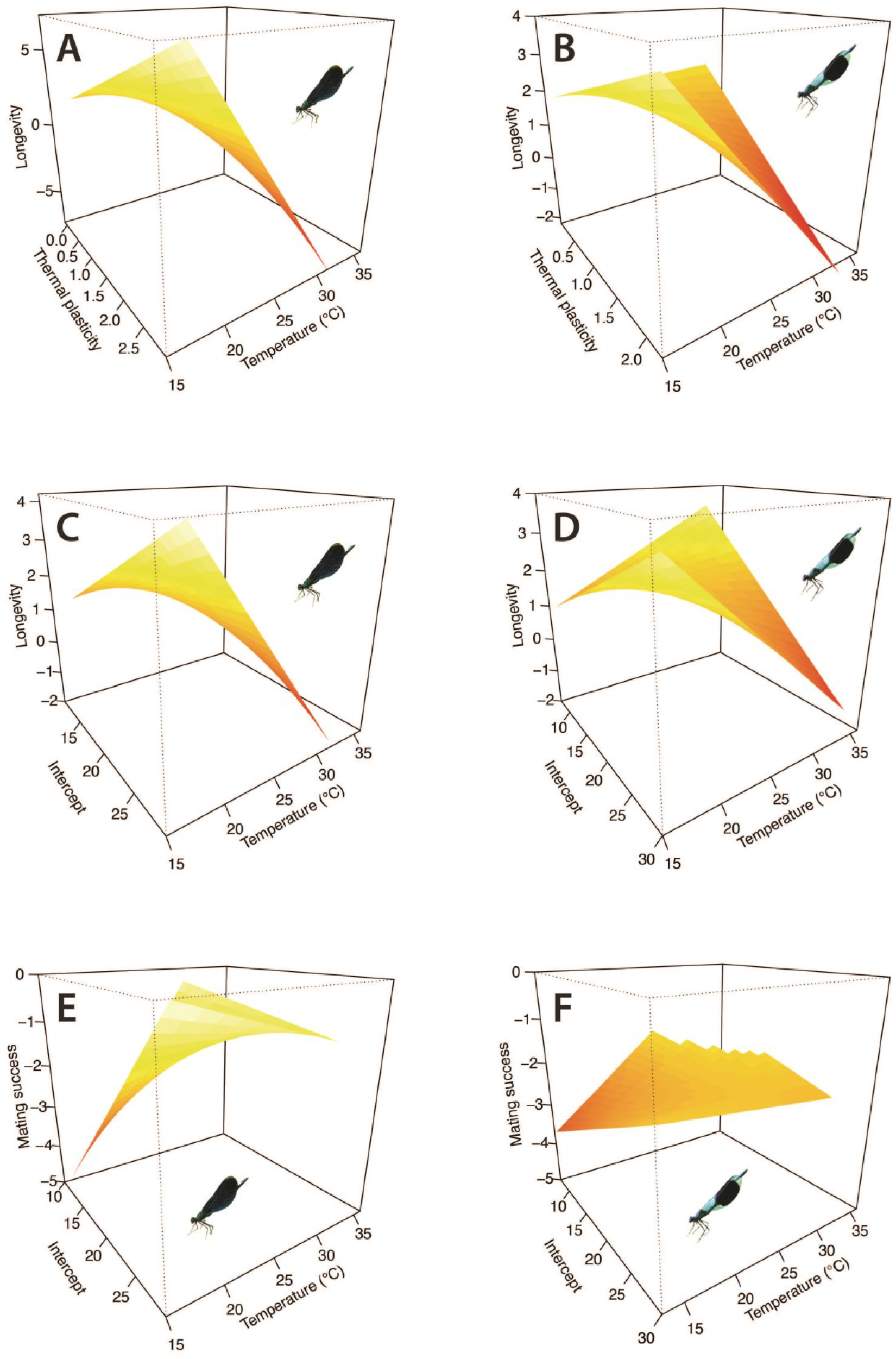
Natural and sexual selection on thermal reaction norm slopes and intercepts in *Calopteryx* males in relation to temperature. Effects of thermal reaction norm parameters (slopes and intercepts; see Fig. 2 and Fig S2) and ambient temperatures on male longevity (**A,C:** *C. virgo,* n = 179; **B, D:** *C. splendens,* n = 778) and for male mating success (**E:** *C. virgo*, n = 413; **F:** *C. splendens*, n=2160). Fitness surfaces (splines) show the results from multivariate generalized additive models (GAM’s)(Table S5). In all four longevity subplots (**A-D**), the interactions between ambient temperature and thermal reaction norm slope and intercept are significant and negative (Table S5). This means that at higher temperatures, both lower plasticity (slope) and lower intercepts are favored, suggesting that thermal plasticity leads to overheating which reduces survival in these two ectothermic insects. In contrast, increasing temperature had a positive effect on mating success in both species (Table S5; cf. Fig. 1C, D).

## Discussion

The novel results presented here is one of few empirical examples where the fitness effects of ambient temperature on both males and females and natural and sexual selection on phenotypic plasticity have been quantified in natural populations. We found that the thermal optima for survival and mating success differed, with survivorship generally being highest at lower ambient temperatures, and with mating success increasing with ambient temperature, suggestive of a trade-off between survival and reproduction (Fig. 1; Tables S1-S2). Moreover, the different thermal optima between survival and reproduction were largely concordant between the sexes and across the species. This suggests that this thermally mediated trade-off between survival and reproduction is general, shared between the sexes and accordingly we found no evidence of sexual antagonism (Fig. 1; Tables S1-S2). These results are broadly in agreement with previous studies on other insects, including both other species of damselflies and butterflies, which generally have found that longevity is highest at lower temperature whereas reproductive output increases with higher temperatures (34–39). For ectothermic animals in general, population growth rate seem to peak between 25 and 30 °C (7), which largely coincides with the temperature peak in mating activity for *Calopteryx* (Fig. 1).

Whereas the relationship between mating success and fitness is straightforward in males, the situation in females could be more complex if there are also fitness costs of mating and preceding male mating harassment (40). However, in *Calopteryx* damselflies, female mating success is likely to be strongly correlated with female fecundity and total reproductive output, at least in the temperate zone where oviposition is often constrained by the number of warm and sunny days (37, 38) and short windows of reproductive opportunities in between many rainy and cold days. In the damselfly *Coenagrion puella*, for instance, life-time female egg production increases with 12 eggs for each 1 °C in ambient temperature (38). For these reasons, male and female reproductive fitness increase in a similar fashion with temperature (Fig. 1C-D, G-H), and temperature therefore seems to have concordant effects on the fitness of both sexes. This implies that thermally mediated selection on one sex (e. g. males) could positively influence fitness also for the other sex (e. g. females), consistent with recent research on desert-dwelling *Drosophila* (30). Hence, sexually concordant selection could facilitate climatic adaptation in both males and females. Concordance between male and female fitness in thermally harsh environments can have some important implications for evolutionary rescue by natural and sexual selection, respectively (26, 41, 42). There is increasing interest to link research fields such as thermal adaptation, sex differences, sexual selection and sexual antagonism to achieve a bitter understanding of how these factors interact over various spatial and biogeographic scales and shape climatic adaptation of organisms (30, 43–47).

Our results challenge a common view that phenotypic plasticity is often adaptive, enabling organisms to survive in novel and harsh environments. We found that natural selection on thermal slopes and intercepts became stronger at higher ambient temperatures, when thermal canalization and robustness was instead favored (Fig. 4A-D, Table S5). This suggests that thermal plasticity would become increasingly maladaptive as ambient temperatures increase during the summer months. This should cause some concern about the future persistence of these thermally sensitive species in the face of ongoing climate change. The mechanisms behind such selection against thermal slopes and intercepts could be the risks of overheating, which can lead to several negative physiological consequences and fitness costs, caused by elevated metabolic rates (48) or increased water loss (4, 12, 49).

We underscore that male fertility might be even more sensitive to high temperatures (50) than male mating success and survival, and the female fitness consequences of high temperatures are likely to be similar to the effects on males (Fig. 1). Potentially, natural selection on males could therefore also have a positive effect also on female fitness and contribute to evolutionary rescue, given the lack of any evidence for sexual antagonism in this study (Fig. 1). Therefore, our results might be conservative regarding the total fitness costs of overheating, considering the role of other male fitness component as well as female fitness.. Our data therefore strongly suggest that natural selection favours thermal canalization and increased environmental robustness at high temperatures, and that current level of thermal plasticity is maladaptive. Our conclusions are broadly in agreement with some other recent studies which have found evidence for non-adaptive or even maladaptive plasticity(21–24). The population consequences of natural and sexual selection on males deserves future consideration in studies of evolutionary rescue (14, 42), especially as selection on males has been considered to be of minor importance in the evolution local adaptations, given female demographic dominance (31) and the common view that it is only females who contribute to population growth.

Results in this study call for greater appreciation for how natural and sexual selection can favour thermal canalization and question the efficiency of thermal plasticity as an evolutionary rescue mechanism. The adaptive significance of phenotypic plasticity should therefore not be taken for granted and plastic traits should ideally be subject to as rigorous selection analyses as to those which have been carried out for non-plastic (fixed) phenotypic traits(4, 5). Finally, our results should lead to some concern, as the prospects of evolutionary rescue through selection on phenotypic plasticity(15, 16, 18, 20) followed by genetic assimilation(15) seems limited for these temperature-sensitive insects. Selection in favour of thermal canalization, rather than plasticity, could potentially counteract or delay extinction risk, although this will also depend on the rate of environmental change and the amount of genetic variation in plasticity, among other factors(16, 19, 20). On a more optimistic note, the positive effect of increasing temperatures on reproductive success in these insects (Figs. 1C-D, G-H) might partly compensate for the negative temperature effects on longevity (Figs. 1A-B, E-F), leading towards a thermally-mediated life-history shift with shorter life spans but with elevated reproduction (51).

## Methods

### Field site and general field work procedures

Our field studies were performed at Sövdemölla Gård along the river Klingavälsån in the province of Skåne in southern Sweden (Latitude: 55.601492 N, Longitude: 13.657340 E). We have carried out field studies at this site and along this river since the year 2000. General field work procedures and basic natural history of the study organisms and ecology of this and other study populations are described in detail elsewhere(52–59). The field studies in the present study were carried out during the summers (June and July) of four years (2013 - 2016). In each year, we recorded individual fitness data of males and females (survival and mating success) and obtained thermal performance measures (estimated individual reaction norms) in males of two sympatric damselfly species (*Calopteryx splendens* and *C. virgo*). Fieldwork took place between 10.00 and 15.00 when activity levels were the highest. Damselflies are ectothermic animals which have low activity levels when temperatures are below 15° C and during rainy and windy days (i.e. they do not fly or mate under such conditions)(60). Therefore, we mainly captured, processed, and re-captured individuals on relatively warm and calm days when they were active and visible, and when it was possible to carry out field work.

### Recording longevity and mating success in the field

In conjunction with our recordings of phenotypic and thermal plasticity data, all captured individuals were marked with a unique code with three colors on the last three segments of the abdomen. These unique markings were used to identify individuals and record their subsequent survival (longevity) in the field. Apart from catching and re-capturing single individuals, we also caught as many copulating pairs in the field as possible to obtain estimates of sexual selection through male mating success. We tried to capture as many mating pairs as possible during days with high mating activity, although we could of course not capture all observed pairs. Nevertheless, we recorded all observed copulations in the field (both those that were captured and those that were not) to quantify mating activity (% individuals found mating out of all individuals) during all field days.

### Measuring thermal microenvironmental variation

We placed 30 iButton thermologgers (1-Wire; DS1921G-F5) at different locations within our field site to measure thermal microenvironmental variation. Such temperature loggers are able to record temperature at set intervals for several months. We recorded ambient temperature every hour for the duration of our field study each year. The field site was divided into 10 neighborhoods (sections; three iButtons per neighborhood) along our study site. Neighborhoods were marked with coloured tape, so that when individuals were recaptured, they could always be linked to a particular neighborhood. When ambient temperature is incorporated in our statistical models on male and female longevities, we used temperature data from these temperature loggers, averaged between 10.00 and 15.00 (the main activity period of these damselflies) of all days these individuals were alive. The time period chosen (10.00-15.00) reflects the warmest period during the day, when either too hot or too cold temperatures are influencing the reproductive activities of these damselflies and therefore also when short-term thermal plasticity is therefore presumably of critical importance.

### Quantifying effects of ambient temperatures on longevity and mating success

To quantify the general effect of temperature on male and female longevity in both species, we estimated the relationship between individual longevity (minimum life-span; last day of capture minus first day of capture(52) and the average daytime temperatures experienced by each individual male or female during their lifespans. We also used average daytime temperatures from the day of capture and in models where we estimated how the probability of being found in copula (our measure of mating success: 0 = not copulating; 1 = copulating) was affected by average daytime temperature on the day of capture. We used general linear mixed models (negative binomial for longevity and binomial for mating success) with year as random factor and the main effect of temperature and squared temperatures were incuded as fixed predictor variables, using the four different datasets of the two species (*C. virgo* males and females and *C. splendens* males and females).

### Thermal imaging and quantifying individual variation in thermal reaction norms

A series of thermal images using a thermal image camera (NEC Avio Infrared Technologies H2640)**(27)** were taken of each field-caught individual male. The accuracy of temperature measurements from this thermal image camera is 0.1 °C. From such thermal images, one can estimate body surface temperatures, which is usually highly correlated with internal body temperatures, especially in small ectothermic animals like insects**(33, 61)** (Fig. S1). We quantified individual body temperatures using our previously developed procedures which are described in detail elsewhere**(27)**. The thermal image camera was regularly calibrated during the use in the field against a black body (black plastic lens cover), with emissivity set to 1.0. To quantify thermal reaction norms (heating rates), individual males were first cooled for three minutes in a cooler box at 5° C and thermal equilibrium. They were then placed back into ambient temperature, each within an individual cup, under a thermal imaging camera. Thermal images were then recorded every ten seconds for three minutes (i.e. up to 18 frames per individual, except for those that flew away within 3 minutes). This experiment aimed to simulate how these ectothermic insects heat up with increasing ambient temperature, e.g. after a cold spell or after a night with low temperatures in the early morning hours. Batches of six individuals were cooled and images were taken simultaneously as they warmed up. After the thermal images were taken, individual males were allowed to fly off to the field site and their subsequent survivorship was recorded through recaptures and later used in the selection analyses (see further below). Images were analysed to estimate the body temperature of each individual at each time point, controlling for ambient temperature. We defined thermal plasticity as the slope of the heating rate (° C/second) of individual males after being cooled down in the fridge box. The intercepts (slope elevation) of these reaction norms is the baseline temperature these males had immediately after being released from the cool box, before starting to heat up (Table S3).

### Selection on thermal plasticity in relation to ambient temperatures

We estimated linear selection gradients from models incorporating the two thermal traits (slopes and intercepts of the thermal reaction norms described above). We carried out these analyses for both sexual selection (male mating success) and survival selection (longevity) and subsequently related this to natural variation in ambient temperature. We analyzed the data using general and generalized linear mixed effect models with year as random factor. Univariate linear selection differentials were estimated separately for the slopes and intercepts of thermal reaction norms following the general statistical methodology of Lande and Arnold(62). The two thermal traits (slopes and intercepts of individual thermal reaction norms) were standardized to mean zero and unit variance and our two fitness components (minimum lifespan or male mating success) were divided by the population average (mean fitness) before being used as the response variables in these regression analyses of selection(62). Since significance levels of regressions with Poisson or binomially distributed data can be unreliable, we also carried out complementary generalized mixed effect models using non-relativized fitness components. We used binomial distributions for mating success and Poisson distribution for longevity. We then present the significance levels from these regressions. Apart from estimating selection gradients on thermal slopes and thermal intercepts, we also visualized the fitness surfaces of how these two thermal traits and ambient temperature influenced survival and mating success, using standard procedures(63).

In the sexual selection analyses, we used a binary response variable as our measure of male mating success (males that obtained at least one mating = 1; non-mated males = 0). From daily field counts and captures at our study site, we found 902 males mating, out of a total of 5902 observed (902/5902 = 0.152). Based on these numbers, we conservatively assume that 15% of males in this population successfully mated at least once. This estimate is likely to be somewhat conservative, as we certainly were not able to record all matings in the field, e. g. those that took place when we did not scan the stream. We used our conservative measure of average male mating fitness (i.e. 0.15) in our regression analyses of selection. This low mating success estimate of 15% males obtaining at least one copulation at this site is consistent with similar low estimates from our previous independent selection studies at two other sites (“Klingavälsåns Naturreservat” and “Höje Å, Värpinge”), where we collected similar data from 2001-2003(53, 54). Low mean mating success reflects the reproductive biology of *Calopteryx* damselflies, which are characterized by high mating skew and strong sexual selection. We continued to monitor the population until all of our marked individuals had died. The average lifespan of an adult *C*. *splendens* males in the field is around 3-4 days, although some males can live for more than two weeks (52, 54).

### Statistical procedures

All statistical analyses were carried out using standard functions in the statistical packages lme4 (64), MASS (65) and mcgv (66) in the R statistical environment (67). Code and datasets will be made available after publication at Dryad Digital Repository (https://datadryad.org/). We carried out general and generalized linear mixed models with different error structures, which are specified under each analysis. We include thermal plasticity traits (slopes and intercepts), ambient temperature and its squared component as fixed predictor variables (to account for curvilinear relationships; see Fig. 1) and year as random effect. We visualized the fitness surfaces (Fig. 4) using splines obtained from Generalized Additive Models (GAM:s)(63).

## Supporting Information

**Table S1. Effects of ambient temperatures on fitness components in male and female *Calopteryx* damselflies.**

**Table S2. Comparisons (within sexes and between species) of the effects of ambient temperatures on fitness components in male and female *Calopteryx* damselflies.**

**Table S3. Species differences in thermal slopes and intercepts of male *Calopteryx*.**

**Table S4. Natural and sexual selection on thermal reaction norm slopes and intercepts in *Calopteryx* males across all seasons and temperatures.**

**Table S5. Natural and sexual selection on thermal reaction norm slopes and intercepts in *Calopteryx* males in relation to ambient temperature.**

**Fig. S1. Fig. S1. External and internal body temperature recordings of individual *Calopteryx* males.**

**Fig. S2. Individual and species variation in thermal reaction norms of male *Calopteryx* damselflies.**

## Acknowledgements

We are grateful to the many field assistants, PhD-students and postdocs who have participated in the field work and thereby directly and indirectly contributed to this study over many years since it started in 2013. We thank Simon Tye for help with the visualizations of Fig. 4. Funding for this study have been provided by research grants from The Swedish Research Council (VR: grant no. 2016-03356), Carl Tryggers Foundation (CTS), Gyllenstiernska Krapperupstiftelsen (grant no. KR2018-0038), Lunds Djurskyddsfond, The Royal Physiographic Society in Lund, Stiftelsen Olle Engqvist Byggmästare and Stina Werners Foundation. We are grateful to the late Jonathan “Jackie” Brown, Adam Hasik, Andreas Nord and Adam Siepielski for critical feedback and comments on the first draft of this manuscript.

## Author Contributions

E.I.S. conceived the idea of this study, obtained funding and planned the field experiments in collaboration with J.W. and M.G.L. J.W., M.G.L. and E.I.S. carried out the field studies and experiments under the guidance from E.I.S. J.W. and M.G.L. performed the statistical analyses with input from E.I.S. E.I.S. wrote the paper. All authors contributed to the finalization of the manuscript.

## Author Information

The authors declare no competing financial interests. Correspondence and requests for materials should be directed to E.I.S. Original data behind all the analyses in this paper and associated R-code will be uploaded on Dryad (https://datadryad.org/).

