## Supporting Material for "Selection on phenotypic plasticity favors thermal canalization"

**This PDF file includes:**

Figs. S1 to S2
Tables S1 to S5

**Table S1. Effects of ambient temperatures on fitness components in male and female *Calopteryx* damselflies.** Statistical analyses below come from generalized linear mixed models and refer to the results in Fig. 1**A-H**. We analyzed temperature effect on male (**A** and **B**) and female survival (**E** and **F**), as well as mating success in natural populations in males (**C** and **D**) and females (**G** and **H**) for *C. splendens* and *C. virgo*.

**A. Male longevity**

*C. virgo* (Fig. 1A)

| Effect | Estimate (SE) | z | *P* |
|---|---|---|---|
| Intercept | **1.678 (0.14)** | **11.31** | **<0.001** |
| Temperature | **2.099 (0.44)** | **4.73** | **<0.001** |
| Temp$^2$ | **-2.371 (0.44)** | **-5.27** | **<0.001** |

**Random effect $\sigma^2$ = 0.061 sd = 0.24**

*C. splendens* (Fig. 1B)

| Effect | Estimate (SE) | z | *P* |
|---|---|---|---|
| Intercept | **1.531 (0.05)** | **27.42** | **<0.001** |
| Temperature | **2.159 (0.22)** | **9.62** | **<0.001** |
| Temp$^2$ | **-2.319 (0.22)** | **-10.20** | **<0.001** |

**Random effect $\sigma^2$ = 0.008 sd = 0.08**

**B. Male mating success**

*C.virgo* (Fig. 1C)

| Effect | Estimate (SE) | z | *P* |
|---|---|---|---|
| Intercept | **-1.784 (0.23)** | **-7.70** | **<0.001** |
| Temperature | **3.631 (1.003)** | **3.62** | **<0.001** |
| Temp$^2$ | **-3.224 (0.96)** | **-3.33** | **<0.001** |

**Random effect $\sigma^2$ = 0.131 sd = 0.36**

*C. splendens* (Fig. 1D)

| Effect | Estimate (SE) | z | *P* |
|---|---|---|---|
| Intercept | **-2.367 (0.16)** | **-14.73** | **<0.001** |
| Temperature | **2.028 (0.53)** | **3.76** | **<0.001** |
| Temp$^2$ | **-1.663 (0.50)** | **-3.26** | **<0.001** |

**Random effect $\sigma^2$ = 0.066 sd = 0.25**

**C. Female longevity**

*C.virgo* (Fig. 1E)

| Effect | Estimate (SE) | z | *P* |
|---|---|---|---|
| Intercept | **1.544 (0.16)** | **9.45** | **<0.001** |
| Temperature | 2.864 (1.63) | 1.75 | 0.079 |
| Temp$^2$ | **-3.245 (1.63)** | **-1.98** | **0.046** |

**Random effect $\sigma^2$ = 0.021 sd = 0.14**

*C. splendens* (Fig. 1F)

| Effect | Estimate (SE) | z | *P* |
|---|---|---|---|
| Intercept | **1.755 (0.33)** | **5.16** | **<0.001** |
| Temperature | **3.364 (1.02)** | **3.28** | **0.001** |
| Temp$^2$ | **-3.771 (1.03)** | **-3.65** | **<0.001** |

**Random effect $\sigma^2$ = 0.315 sd = 0.56**

**D. Female mating success**

*C.virgo* (Fig. 1G)

| Effect | Estimate (SE) | z | *P* |
|---|---|---|---|
| Intercept | -0.155 (0.48) | -0.31 | 0.751 |
| Temperature | -0.841 (1.20) | -0.69 | 0.487 |
| Temp$^2$ | 0.784 (1.18) | 0.66 | 0508 |

**Random effect $\sigma^2$ = 0.657 sd = 0.81**

*C. splendens* (Fig. 1H)

| Effect | Estimate (SE) | z | *P* |
|---|---|---|---|
| Intercept | **-0.676 (0.31)** | **-2.13** | **0.032** |
| Temperature | **3.297 (0.78)** | **4.17** | **<0.001** |
| Temp$^2$ | **-2.902 (0.76)** | **-3.81** | **<0.001** |

**Random effect $\sigma^2$ = 0.279 sd = 0.52**

In these mixed models, Year was included as a random factor to control for intercept variation between the three different seasons (2014-2016). Standardized temperature and squared temperature were included as fixed (continuous) predictor variables in these models, standardization was achieved by subtracting the mean to each temperature value and dividing by the standard deviation. We run a separate analysis for each of the four phenotypic categories (males and females of each species) and fitness components (survival and mating success). In the survival analyses, we treated longevity as response variable in a negative binomial model to account for overdispersion of the data. Field data on mating success were analyzed using a binomial distribution for the dependent fitness variable (0=unmated; 1=mated). Significant effects are marked in bold.

**Table S2. Comparisons (within sexes and between species) of the effects of ambient temperatures on fitness components in male and female *Calopteryx* damselflies.** Statistical analyses below come from generalized linear mixed models and refer to the results in Fig. 1**A-H**. Each model comes from one row within Fig. 1, comparing one phenotypic category (females or males) and one fitness component (survival or mating success)(**A** vs. **B**, **C** vs. **D**, **E** vs. **F** and **G** vs. **H**).

**Male longevity (Fig. 1A vs. B)**

| Effect | Estimate (SE) | z | *P* |
|---|---|---|---|
| Species (*C. virgo*) | **0.119 (0.04)** | **2.78** | **0.005** |
| Temperature | **2.160 (0.07)** | **9.80** | **< 0.001** |
| Temperature^2 | **-2.360 (0.002)** | **-10.55** | **< 0.001** |
| Species (*C. virgo*) : Temperature | 0.005 (0.52) | 0.18 | 0.85 |
| Species (*C. virgo*) : Temperature^2 | -0.077 (0.52) | -0.14 | 0.88 |

N=2412

**Male mating success (Fig. 1C vs. D)**

| Effect | Estimate (SE) | z | *P* |
|---|---|---|---|
| Species (*C. virgo*) | 0.440 (0.12) | 3.51 | **<0.001** |
| Temperature | **2.263 (0.55)** | **4.10** | **<0.001** |
| Temperature^2 | **-1.834 (0.53)** | **-3. 46** | **<0.001** |
| Species (*C. virgo*) : Temperature | 0.722 (1.07) | 0.67 | 0.502 |
| Species (*C. virgo*) : Temperature^2 | -0.811 (1.01) | -0.79 | 0.425 |

N=4810

**Female longevity (Fig. 1E vs. F)**

| Effect | Estimate (SE) | z | *P* |
|---|---|---|---|
| Species (*C. virgo*) | -0.144 (0.18) | -0.80 | 0.42 |
| Temperature | **3.546 (0.96)** | **3.68** | **<0.001** |
| Temperature^2 | **-3.844 (0.97)** | **-3. 59** | **< 0.001** |
| Species (*C. virgo*) : Temperature | -0.322 (2.24) | -0.14 | 0.88 |
| Species (*C. virgo*) : Temperature^2 | 0.094 (2.23) | 0.04 | 0.96 |

N=159

**Female mating success (Fig. 1G vs. H)**

| Effect | Estimate (SE) | z | *P* |
|---|---|---|---|
| Species (*C. virgo*) | **0.457 (0.16)** | **2.74** | **0.006** |
| Temperature | **3.519 (0.78)** | **4.46** | **<0.001** |
| Temperature^2 | **-3.086 (0.76)** | **-4.01** | **<0.001** |
| Species (*C. virgo*) : Temperature | **-4.600 (1.50)** | **-3.06** | **0.002** |
| Species (*C. virgo*) : Temperature^2 | **3.964 (1.43)** | **2.75** | **0.005** |

N=1090

**Table S3. Species differences in thermal slopes and intercepts of male *Calopteryx*.** We used t-tests to compare species-differences in the slope (Fig. 2B) and intercept (Fig. 2C) of male thermal reaction norms (Fig. S2).

| Thermal reaction norm trait | *C. virgo* (mean ± SD) | *C. splendens* (mean ± SD) | *P* |
|---|---|---|---|
| Intercept (Initial temperature: °C) | **19.238 (± 3.18)** | **17.364 (3.17)** | **<0.001** |
| Slope (Thermal plasticity: °C/s) | 0.534 (0.34) | 0.508 ( ± 0.26) | 0.339 |

N=957. Significant effects are marked in bold.

**Table S4. Natural and sexual selection on thermal reaction norm slopes and intercepts in *Calopteryx* males across all seasons and temperatures**. To estimate selection gradients on thermal reaction norm slopes and intercepts we used linear mixed models (see section "Selection on thermal plasticity in relation to ambient temperatures"). Significance levels were obtained using generalized linear mixed models assuming binomial (for mating success) and a Poisson distribution (for longevity). For clarity and brevity, we do not include the parameter value of the intercept in the four models presented here.

| | *C. splendens* longevity (Fig. 3A) | | | *C. splendens* mating success (Fig. 3A) | | |
|---|---|---|---|---|---|---|
| Effect | Estimate (SE) | *t* | *P* | Estimate (SE) | *t* | *P* |
| Slope | **-0.088 (0.04)** | **-2.193** | **< 0.001** | 0.051 (0.06) | 0.832 | 0.418 |
| Intercept | **-0.077 (0.04)** | **-1.857** | **< 0.001** | **0.171 (0.06)** | **2.571** | **0.011** |
| N=778 | | | | N=2165 | | |

| | *C. virgo* longevity (Fig. 3B) | | | *C. virgo* mating success (Fig. 3B) | | |
|---|---|---|---|---|---|---|
| Effect | Estimate (SE) | *t* | *P* | Estimate (SE*)* | *t* | *P* |
| Slope | **0.070 (0.07)** | **0.995** | **0.012** | -0.133 (0.11) | -1.199 | 0.230 |
| Intercept | **0.134 (0.07)** | **1.885** | **0.002** | **0.404 (0.11)** | **3.488** | **0.001** |
| N=179. | | | | N=413 | | |

Significant effects are marked in bold.

**Comparing male selection differentials for longevity (*C. virgo* vs. *C. splendens*; Fig. 3A vs. 3B)**

| Effect | Estimate (SE) | z | *P* |
|---|---|---|---|
| Species (*C. virgo*) | -0.073 (0.09) | -0.51 | 0.94 |
| Slope | **-0.093 (0.04)** | **-5.45** | **<0.001** |
| Intercept | **-0.084 (0.04)** | **-6.06** | **<0.001** |
| Species (*C. virgo*):Slope | **0.157 (0.07)** | **5.74** | **<0.001** |
| Species (*C. virgo*):Intercept | **0.198 (0.09)** | **5.87** | **<0.001** |
| N=957 | | | |

**Comparing male selection differentials for mating success (*C. virgo* vs. *C. splendens*; Fig. 3A vs. 3B)**

| Effect | Estimate (SE) | z | *P* |
|---|---|---|---|
| Species (*C. virgo*) | -0.185 (0.15) | 1.10 | 0.27 |
| Slope | 0.044 (0.05) | 0.62 | 0.52 |
| Intercept | **0.194 (0.06)** | **2.91** | **0.003** |
| Species (*C. virgo*):Slope | -0.152 (0.14) | -1.25 | 0.21 |
| Species (*C. virgo*):Intercept | 0.139 (0.14) | 1.35 | 0.18 |
| N=2578 | | | |

**Table S5. Natural and sexual selection on thermal reaction norm slopes and intercepts in *Calopteryx* males in relation to ambient temperature.** Statistical tests of the response surfaces in Fig. 4. To analyze the effects of slope and intercept in relation to ambient temperature and their effects on longevity (natural selection) and mating success (sexual selection) we used Generalized Additive Models (GAMs). In all these models we standardized slopes and intercepts to mean zero and unit variance, and included main effects of slope, intercept and temperature and all the three two-way interactions (slope x intercept, slope x temperature and intercept x temperature). We used a negative binomial and a binomial distribution for longevity and mating success, respectively. For clarity, we omitted the model intercepts of all four models here (*P* <0.001 in all cases), to avoid confusion with the entirely different thermal trait intercept for which we estimated phenotypic selection (Fig. 4A-F).

| | ***C. splendens* longevity (Fig. 4B,D)** | | | ***C. splendens* mating success(Fig. 4F)** | | |
|---|---|---|---|---|---|---|
| Effect | Estimate (SE) | z | *P* | Estimate (SE) | z | *P* |
| Slope | **-0.095 (0.03)** | **-2.844** | **<0.001** | -0.004 (0.08) | -0.079 | 0.955 |
| Temperature | **-0.148 (0.03)** | **-4.363** | **<0.001** | **0.249 (0.09)** | **2.749** | **0.005** |
| Intercept | -0.017 (0.03) | -0.540 | 0.589 | 0.045 (0.08) | 0.511 | 0.609 |
| Slope x Temperature | **-0.134 (0.03)** | **-3.891** | **<0.001** | 0.033 (0.08) | 0.406 | 0.685 |
| Temperature x Intercept | **-0.167 (0.03)** | **5.238** | **<0.001** | -0.080 (0.07) | -1.112 | 0.266 |
| Slope x Intercept | 0.059 (0.03) | 0.036 | 0.106 | 0.030 (0.08) | 0.388 | 0.698 |
| N=778. | | | | N=2165. | | |

| | ***C. virgo* longevity (Fig. 4A,C)** | | | ***C. virgo* mating success (Fig. 4E)** | | |
|---|---|---|---|---|---|---|
| Effect | Estimate (SE) | z | *P* | Estimate (SE) | z | *P* |
| Slope | 0.006 (0.06) | 0.098 | 0.921 | -0.327 (0.18) | -1.736 | 0.082 |
| Temperature | **-0.478 (0.06)** | **-7.307** | **<0.001** | **0.437 (0.18)** | **2.406** | **0.016** |
| Intercept | 0.106 (0.06) | 1.765 | 0.077 | 0.312 (0.18) | 1.702 | 0.088 |
| Slope x Temperature | **-0.295 (0.08)** | **-3.341** | **<0.001** | -0.183 (0.19) | -0.941 | 0.346 |
| Temperature x Intercept | **-0.205 (0.06)** | **-3.036** | **0.002** | -0.195 (0.15) | -1.229 | 0.219 |
| Slope x Intercept | 0.063 (0.04) | 1.339 | 0.180 | -0.275 (0.14) | -1.956 | 0.050 |
| N=179. | | | | N=413. | | |

Significant effects are marked in bold.

**Legends to Supporting Figures**

**Fig. S1. External and internal body temperature recordings of individual *Calopteryx* males.** We recorded individual external (grey) temperature using thermal imaging, and simultaneously the internal body temperature (black) using a copper thermal thermometer coupled to a cable extension and inserted inside the damselfly thorax (n=7 males). A generalized linear mixed model using temperature type (external or internal) and time point as fixed factor, and individual male as random effect, showed no difference in the slope of body temperature between external and internal measurements (estimate = -0.011, std. error = 0.03, p = 0.757). Shown are individual temperature estimates (data points from individual males) and change in temperature with time (X-axis), estimated in a similar fashion as the thermal reaction norms (Fig. 2A; Fig. S2).

**Fig. S2. Individual and species variation in thermal reaction norms of male *Calopteryx* damselflies.** Individual variation in thermal reaction norms of **A.** *C. splendens* (left) and **B.** *C. virgo* (right). Thick solid lines represent the average thermal reaction norm in the population, Y-axis thorax temperature (in °C) and X-axis time (in seconds) from the time point when the individual male was taken out of the cooler. The two species have similar slopes of their thermal reaction norms (Fig. 2B; Table S2), but they differ significantly in their intercepts (elevations), with *C. virgo* having higher baseline thorax temperature than *C. splendens* (Fig. 2C; Table S3).

**Fig. S1**

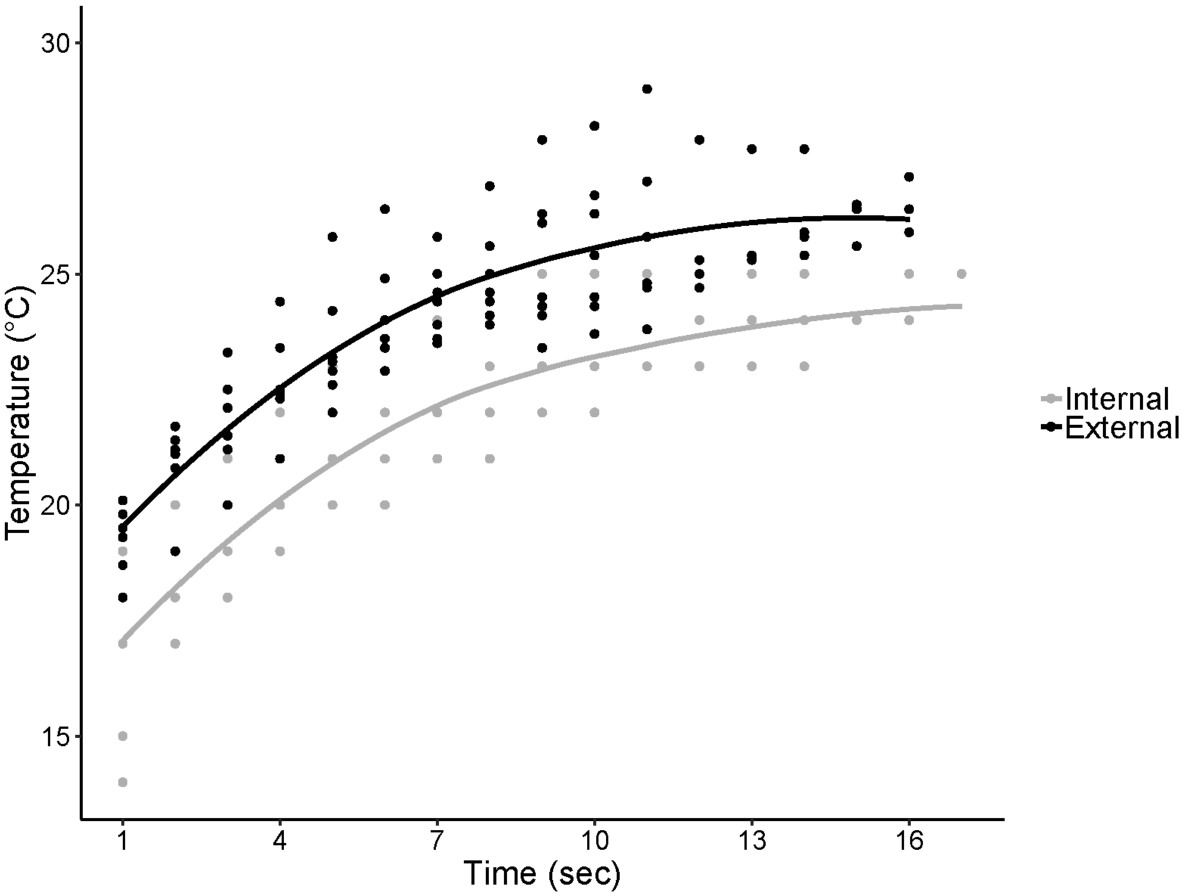

30
25
20
15
Temperature (°C)
1
4
7
10
13
16
Time (sec)
Internal
External

**Fig. S2**

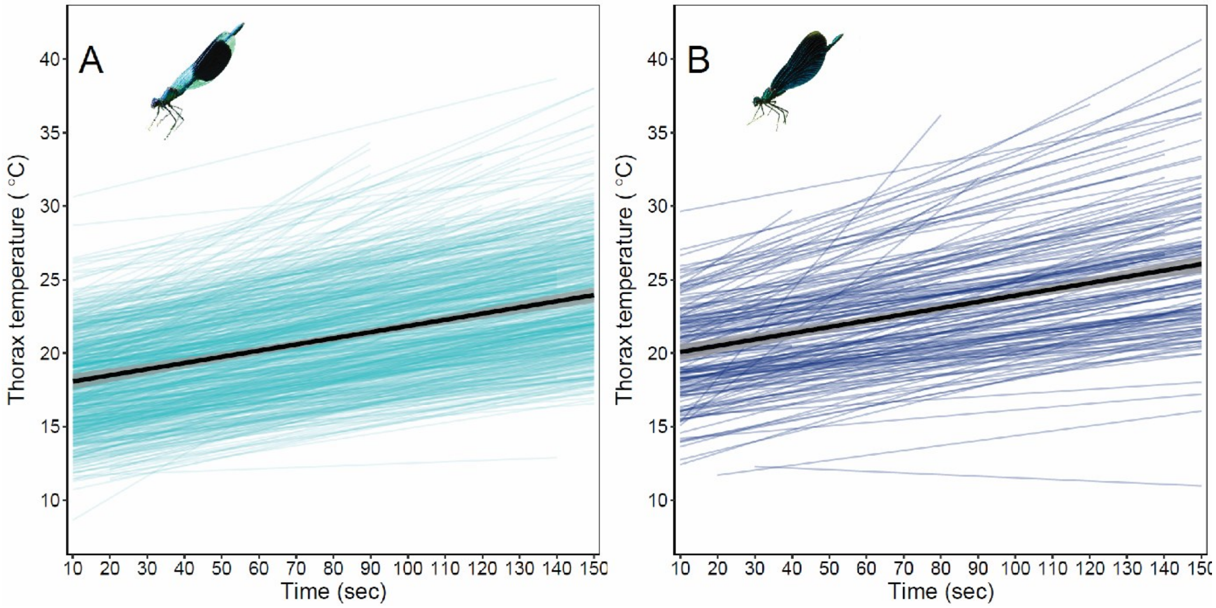
A
B
Thorax temperature ( °C)
Time (sec)
10
15
20
25
30
35
40
10 20 30 40 50 60 70 80 90 100 110 120 130 140 150
